# ONC206 demonstrates potent anti-tumorigenic activity and is a potential novel therapeutic strategy for high-risk medulloblastoma

**DOI:** 10.1101/2025.09.25.678693

**Authors:** Theophilos Tzaridis, Jingbo Liu, Franklin Lee Chien, Anshu Malhotra, Dan Zhu, Isabella Gershon, Hongying Zhang, Jose E. Velazquez Vega, Matthew Schniederjan, Teresa Sposito, Peter D. Adams, Joshua E. Allen, Varun V. Prabhu, Robert J. Wechsler-Reya, Tobey J. MacDonald

**Affiliations:** Cancer Genome and Epigenetics Program, NCI-Designated Cancer Center, Sanford Burnham Prebys Medical Discovery Institute, La Jolla, CA, USA; Department of Pediatrics, Emory University School of Medicine, Atlanta, GA, USA; Emory University, Atlanta, GA USA; Department of Pathology, Children’s Healthcare of Atlanta, Atlanta, GA, USA; Division of Regenerative Medicine, Department of Medicine, Moores Cancer Center and Sanford Stem Cell Institute, UC San Diego, La Jolla, CA, USA; Chimerix (a Jazz Pharmaceuticals Company), Durham, NC; Department of Neurology and Herbert Irving Comprehensive Cancer Center, Columbia University Medical Center, New York, NY, USA; Aflac Cancer and Blood Disorders Center, Children’s Healthcare of Atlanta, Atlanta, GA, USA

**Keywords:** Medulloblastoma, novel therapy, ONC206, mitochondrial damage, ClpP

## Abstract

**Background:** Medulloblastoma is the most common malignant pediatric brain tumor, and has an urgent need for novel treatment approaches. Dordaviprone (ONC201) and its chemical derivative with nanomolar potency, ONC206, induce apoptosis of cancer cells by activation of the mitochondrial caseinolytic protease P (ClpP). ONC206 is currently in Phase I clinical trials for pediatric patients with primary brain tumors.

**Methods:** In this study, we evaluated the preclinical therapeutic effects of ONC206 in medulloblastoma and investigated its mechanism of action.

**Results:** We found evidence for high expression of ClpP at both the RNA and protein level in medulloblastoma tumors, compared to very low expression in normal brain tissue. In addition, we saw a pronounced reduction in cell viability of human Group 3 and Group 4 and murine SHH-driven and Group 3 medulloblastoma cells treated with ONC206 with low IC-50s. After treatment with ONC206, we observed an induction of integrated stress response and mitochondrial damage. To test the efficacy of ONC206 *in vivo*, we used murine models of SHH-driven and Group 3 medulloblastoma as well as Group 3 and Group 4 patient-derived xenografts (PDXs). ONC206 led to a significant prolongation of survival in both murine models, with the SHH mice demonstrating survival extension from 70 to 140 days. PDX-bearing mice also responded to ONC206, which led to a significant survival benefit.

**Conclusion:** Our results highlight ONC206 as a novel therapeutic option for patients with high-risk medulloblastoma and provide strong rationale for testing the efficacy of ONC206 in the treatment of these patients.

**Key points (2-3):** ONC206 potently kills medulloblastoma cells by inducing integrated stress response and mitochondrial damage.

ONC206 prolongs survival of medulloblastoma-bearing mice in both murine and patient-derived xenograft models.

**Importance of study:** There is an unmet need for better therapies for high-risk medulloblastoma patients. ONC201 has shown promising responses and recently received FDA approval for diffuse midline glioma. ONC206 is a chemical derivative with higher potency and better brain penetrance. In this study, we analyzed the therapeutic potential of ONC206 for high-risk medulloblastoma and found that the drug effectively killed mouse and human medulloblastoma cells with high nanomolar potency. We also saw that ONC206 very significantly prolonged survival of medulloblastoma-bearing mice, both in genetically engineered mouse models and patient-derived xenografts. Our study provides a strong rationale for testing the efficacy of ONC206 in the treatment of patients with medulloblastoma and has set the stage for a clinical trial with this agent in pediatric patients with recurrent malignant brain tumors, including medulloblastoma (NCT04732065).

## Introduction

Medulloblastoma (MB), the most common pediatric malignant brain tumor, accounts for approximately 20% of all pediatric cancers and 70% of all embryonal tumors of the central nervous system (CNS), and is associated with a high mortality rate. ^1,2^ MB consists of four molecular and clinicopathological prognostic subgroups – Wingless (WNT)-activated, Sonic-

Hedgehog-activated (SHH), Group 3 and Group 4,^3^ which have been further refined in the most recent WHO classification of 2021.^4^ Despite considerable improvement in the 5-year overall survival (OS) rate to more than 80% for average-risk disease, children with high-risk disease continue to have relatively poor OS (∼70%), and in those with high-risk SHH or metastatic *Myc*-amplified Group 3 tumors, OS remains dismal (<55%) ^5^. Current standard of care treatment comprises surgical resection, cranio-spinal irradiation (CSI) in non-infants, and intensive multimodal chemotherapy; however, this treatment frequently results in long-term neurological and endocrine sequelae ^6^. Moreover, recurrent MB has limited therapeutic options and is frequently fatal. Therefore, novel targeted therapies improving not only survival, but also quality of life for patients with this devastating disease are urgently needed.

Imipridones are novel agents shown to selectively kill cancer cells, while being much less toxic to normal cells ^7^. Interestingly, while initially believed to function mainly through inhibition of the dopamine receptor D2 (DRD2) ^8,9^ or induction of tumor necrosis factor (TNF)-related apoptosis-inducing ligand (TRAIL) signaling ^10,11^, recent data have suggested that they exert their function through activation of the mitochondrial caseinolytic protease P (ClpP), which leads to selective mitochondrial protein degradation and thereby *TP53* independent apoptosis of tumor cells ^12,13^. One prominent example of imipridones is ONC201, which has demonstrated preclinical activity, as well as brain penetrance, safety and clinical responses in patients with recurrent glioblastoma ^14,15^. Based on those encouraging results, additional trials were initiated in pediatric and adult patients with *H3K27M* mutant diffuse midline gliomas (DMG) and showed very encouraging results with sustained tumor responses in a subset of these patients ^16,17^. Importantly, ONC201 is now the first FDA-approved drug for the treatment of patients with progressive DMG. While ONC201 demonstrates cytostatic effects at a micromolar range, its derivative ONC206 is effective at nanomolar concentrations, shows a stronger reduction of ATP levels compared to ONC201 and has similar oral bioavailability and tolerability ^18^. Notably, ONC206 is currently under investigation for the treatment of patients with recurrent DMG and other refractory primary malignant brain tumors (NCT04541082 and NCT04732065), and prior reports have described a dependency on mitochondrial bioenergetics for maintaining medulloblastoma survival and proliferation ^19–21^.

In this study, we evaluated the molecular mechanism of action and preclinical therapeutic effects of ONC206 for the treatment of medulloblastoma using *in vitro* assays and both human and murine *in vivo* models of MB. Herein, we report high on-target efficacy, as well as good tolerability, of ONC206 for the treatment of genetically engineered murine models of SHH-driven and Group 3 MB in addition to Group 3 and Group 4 patient derived xenograft (PDX) MB models.

## Materials and Methods

### Molecular Data Mining

We interrogated publicly available databases to mine the data for gene expression of ClpP and DRD2 in medulloblastoma, normal brain and normal cerebellum tissue (R2: Medulloblastoma Genomics Analysis and Visualization Platform, https://hgserver1.amc.nl/cgi-bin/r2/main.cgi?dscope=MB&option=main and NCBI Gene Expression Omnibus (GEO) GEO2R; https://www.ncbi.nlm.nih.gov/geo/query/acc.cgi?acc=GSE85217), and protein expression of ClpP in medulloblastoma relative to other pediatric brain tumors and normal cerebellum tissue (CPTAC Data Portal https://proteomics.cancer.gov/data-portal). The p-values and expression data plots were generated by log-rank tests using the analytical and graphical software available through the respective website-based databases.

### Immunohistochemistry

Tumor and normal cerebellum tissues were fixed in 10% neutral buffered formalin, processed, and embedded in paraffin. Five micrometer thick sections were deparaffinized in xylene and rehydrated to water prior to microwave antigen retrieval in Tris/HCl/EDTA pH 9.0 buffer (Dako Cytomation, Glostrup, Denmark) and PBS washing. After neutralization of the endogenous peroxidase with 3% H2O2 for 10 min, the sections were incubated with protein blocking buffer for 10 min before undergoing incubation with the primary antibody: Cleaved Caspase3 (PA5-77887, Thermo Fisher Scientific), ClpP (ab8416, ABCAM), C-Myc (#13987, Cell Signaling), DRD2 (#22022-1-AP, Thermo Fisher Scientific), DR5 (ab8416, ABCAM), EZH2 (#5246, Cell Signaling), NDUFS7 (NBP1-49846, Novus Biologicals), and TGM2 (#3557, Cell Signaling). Staining was developed using DAB (Vector Laboratories, Burlingame, CA, USA) followed by Hematoxylin counterstaining (Millipore Sigma). H&E staining was also performed to access the location of the tumor.

### Cell culture

Established cell lines (D556, Daoy and UW-228) were obtained from ATCC (Manassas, VA, USA) and cultured in Dulbecco’s Modified Eagle Medium (DMEM) supplemented with 10% fetal bovine serum (all ThermoFisher, Waltham, MA, USA).

PDX cells were grown in immunocompromised mice (strain details below) and once harvested (see protocol below), they were cultured in suspension flasks (Genesee Scientific, El Cajon, CA, USA) with Neurobasal-A medium containing supplements N2 and serum-free B27, as well as GlutaMAX (100U/ml), Penicillin-Streptomycin (100U/ml) and Heparin (2μg/ml, all ThermoFisher Scientific).

Cells were regularly tested for mycoplasma via polymerase chain reaction (PCR).

### Western Blot

Medulloblastoma cells were washed and harvested in PBS and lysates were prepared as previously described ^22^. 30µg of total protein each was separated by SDS-PAGE. Western Blotting was carried out using standard methods. The primary antibodies used were ATF4, ClpP, DR5, GAPDH, LC3-b, MitoProfile total OXPHOS antibody, SDHA, Survivin and Tomm20 (ABCAM); eIF2a, pS473-Akt, STAT3 and pY705-STAT3 (Cell Signaling). HRP-conjugated secondary antibodies were anti-mouse (Jackson Immuno Research) or anti-rabbit (Pierce, Life Technologies).

### Real time quantitative RT PCR

Total RNA was prepared from the mice cerebellum tissue. Random primed single stranded cDNA was made from total RNA by using the Superscript III kit (Cat# 18080 051; Thermo Fisher Scientific). The following oligonucleotides as primers were used for real time PCR: murine Glyceraldehyde 3 phosphate dehydrogenase (GAPDH), 5′ CATCACTGCCACCCAGAAGACTG 3′ (forward), 5′ ATGCCAGTGAGCTTCCCGTTCAG 3′ (reverse); murine C-Myc, 5′ TCGCTGCTGTCCTCCGAGTCC 3′ (forward), 5′ GGTTTGCCTCTTCTCCACAGAC 3′ (reverse); murine Ezh2, 5′ CATACGCTCTTCTGTCGACGATG 3′ (forward), 5′ ACACTGTGGTCCACAAGGCTTG 3′ (reverse); murine DR5, 5′ GCAGAGAGGGTATTGACTACACC 3′ (forward), 5′ GCATCGGGTTTCTACGACTTT 3′ (reverse). Data analysis was performed according to the absolute standard curve method. Data are presented in relation to the respective housekeeping gene and normalized the fold change of control cells to 1 (100%), then calculated the relative fold changes.

### Mitochondrial assays

Medulloblastoma cells were seeded at low density on 1mm glass coverslips and cultured for 48 h in DMEM containing 10% FBS. Cells were then incubated with 100 nM MitoTracker Red probe (Invitrogen) for 30 minutes according to manufacturer’s instructions. Immunofluorescence staining was carried out following standard protocols. Cells were fixed with 4% paraformaldehyde in PBS for 10 min and then washed with PBS and permeabilized with 0.1% Triton X-100 for 5 min and washed again with PBS. After fixation, glass coverslips were incubated with primary anti-Cytochrome c antibody (Cell Signaling) and then secondary FITC-labeled anti-mouse antibody (Thermo Fisher Scientific) or 300 nM DAPI nuclear stain (Cell Signaling). The samples were then mounted on microscope slides in ProLong® Gold Antifade Mountant (Life Technologies). Images were acquired immediately thereafter. Z-stack images were acquired using an Olympus FV1000 confocal microscope.

### Cell viability assay

Cells were seeded in 96-well plates (50,000 cells per well) and were treated for 72 hours with different concentrations of ONC201 and ONC206, a DMSO control was included. After treatment, cell viability was determined using CellTiter-Glo (Promega, Madison, USA). For IC-50 determination, a non-linear regression analysis was applied (GraphPad Prism, La Jolla, USA).

### In vivo experiments

#### ONC206 treatment of SmoA1-GFP murine transgenic model of SHH-type MB

ND2:SmoA1 transgenic mice that spontaneously form SHH-driven murine MB tumors were purchased from The Jackson Laboratories (Bar Harbor, ME, USA). To generate ND2:SmoA1-GFP mice for this study, homozygous ND2:SmoA1 mice (SmoA1) were crossed with Math1-GFP reporter mice kindly provided by Dr. Tracy-Ann Read, Emory Winship Cancer Institute. The F1 (SmoA1–Math-GFP heterozygous) mice were back-crossed with SmoA1 mice. Real-time PCR was used to select the F2 mice homozygous for SmoA1 as well the Math-GFP reporter gene. The selected F2 mice then were back-crossed with SmoA1 for three more generations to yield the final SmoA1-GFP mice. Using fluorescence microscopy, the primary MB tumor of the cerebellum are readily visualized by positive detection of GFP for all tumor localization assessments. All animals were maintained in the animal facility at Emory University and used in accordance with protocols approved by the Emory Institutional Animal Care and Use Committee.

A total of 25 SmoA1-GFP mice with early tumor detection confirmed by imaging by MRI with a 9.4 T Bio-spec scanner (Bruker) at 10 weeks of age were identified for drug treatment studies. For MR imaging, mice were anesthetized with 1.5 to ∼2% isoflurane during the image acquisition. Respiration, body temperature, and electrocardiogram (ECG) were monitored during the imaging. T2-weighted images were acquired using a rapid acquisition with refocused echoes (RARE) sequence in sagittal direction (time to repetition [TR] = 4,000 ms, time to echo [TE] = 42 ms, RARE factor = 8, field of view [FOV] = 20 × 20 mm2, matrix = 116 × 116, thickness = 0.75 mm, 15 slices, and average FOV obtained per image = 12). The tumor volume was qualified as the products of slice-to-slice separation and the sum of areas from the manually drawn tumor regions of interest (ROIs) on images.

The tumor positive mice were randomized into 3 groups of equal age, male: female ratios and equivalent starting tumor volume prior to treatment. One group received oral gavage of ONC206 (120mg/kg ; n=11), the other group (100mg/kg; n=5) and the control group (vehicle control ; n=9) weekly for two weeks. Tumor growth over time by volume was serially measured by MRI. Animals were kept alive until they displayed signs of morbidity or toxicity (>20% weight loss), whereupon they were euthanized.

#### ONC206 treatment of murine Group 3 MB and human Group 3, Group 4 MB PDXs

All animal experiments were approved by the Institutional Animal Care and Use Committee of the Sanford Burnham Prebys Medical Discovery Institute, La Jolla. For experiments with immune-competent mice, female 12-week old B6(Cg)-Tyr^c-2J^/J (Jackson Laboratories, Ben Harbor, USA) were transplanted with previously described *Myc* overexpressing and dominant negative *Trp53* MB cells (20,000 cells per mouse) ^23^. At d8 after transplant, mice were randomized into treatment groups (n=9 per group) based on bioluminescent signaling using in vivo imaging system (IVIS, PerkinElmer, Waltham, USA). Tumor-bearing mice were treated with 100mg/KG ONC206 or vehicle control twice weekly for four weeks. For PDX experiments, 12-week old female athymic nude mice Nu/J (Jackson Laboratories) were transplanted with human MB cells (50-100,000 cells per mouse). At d25-40 after transplant, mice were randomized into treatment groups (n=6-8 per group) and treated similar to our murine MB models. After receiving updates from the NCT04541082 trial on efficacy and tolerability of ONC206, Group 4 RCMB99-bearing mice were treated with 50mg/kg ONC206 or vehicle control twice a day for three days per week until disease-related death. Animals were kept alive until they displayed signs of morbidity or toxicity (>20% weight loss), whereupon they were euthanized.

### Tumor tissue dissociation

Murine tumors were enzymatically dissociated using DNase (2500U/ml, Worthington, Lakewood, NJ, USA) and Liberase (100μg/ml, Sigma-Aldrich, St. Louis, MO, USA). Enzymatic dissociation was blocked using 1% FBS. The suspension was then passed through a 70 micron filter (Corning, Corning, NY, USA) and red blood cell lysis was performed using the appropriate lysis buffer (Invitrogen, Waltham, MA, USA). Single cell suspension was then cultured for *in vitro* treatments based on the protocols mentioned above.

### Statistical analysis

For statistical analyses, GraphPad Prism software version 9 (La Jolla, CA, USA) was used. For IC-50) determination, a non-linear regression analysis was applied. Survival studies were performed using Log-rank tests and survival was visualized via Kaplan Meier curves. Statistical tests used were upaired two-tailed t-tests.

### Methods for Supplementary Data

#### Organotypic slice culture assays

Organotypic slice cultures were prepared as described previously ^24^ using resected human medulloblastoma tumor tissue. The slices were treated directly with ONC206 or control DMSO with or without 5Gy irradiation after 24 hours of treatment with ONC206 or DMSO. Following culture for another 24 hours, slices were fixed with 4% formaldehyde and embedded in OCT medium prior to cryosectioning and then apoptosis of cells was detected using the TUNEL Assay Kit (Cell Signaling) for immunofluorescence microscopy. Z-stack images were acquired using an Olympus FV1000 confocal microscope.

#### ONC206 drug concentration measured in murine brain tumor tissue

Six MRI-confirmed tumor-bearing SmoA1-GFP mice were randomly divided into two group, one received ONC206 by oral gavage at dose of 120mg/kg(n=3), and the other controls (n=3). Mice were euthanized one hour after gavage treatment, tumor tissue and normal cerebellum tissue were collected separately under fluorescence microscopy and sent on dry ice to WuXi AppTec Laboratory Testing Division (6 Cedarbrook Drive, Cranbury, NJ 08512, USA ) for analysis.

<u>Sample Preparation for HPLC:</u> Samples were homogenized in 1:6 (wt:v) ratio using LC-MS water. Samples under 20 mg were homogenized with 1:10 (wt:v) ratio LC-MS water instead. 20 µL sample was protein precipitated with 200 µL IS solution (100 ng/mL Labetalol & 100 ng/mL Tolbutamide in ACN). The mixture was vortex-mixed well and centrifuged at 3900 rpm for 10 min, 4 . An aliquot of 100 μL supernatant was transferred to sample plate and mixed with 100 μL water, then the plate was shaken at 800 rpm for 10 min. and 0.10 µL of supernatant was then injected for LC-MS/MS analysis.

<u>Instrument Conditions:</u> The ONC206 concentration was measured by using SCIEX Triple Quad 6500+ LC/MS on a Waters ACQUITY UPLC® system. Column: Waters ACQUITY UPLC HSS T3 2.1 x 55mm, 1.8µm. Mobile phases A and B comprised 0.1% Formic Acid in water (v/v, aqueous) and 0.1% Formic Acid in ACN (v/v, organic), respectively at a flow rate of 0.6 mL/minute. The product ions of ONC-206: [M+H]+ m/z 409.2>, Labetalol(IS): [M+H]+ m/z 329.2>162.1and Tolbutamide(IS): [M+H]+ m/z 271.1>155.3 were detected via multiple reaction monitoring(MRM) on mass spectrometer in ESI positive mode.

## Results

### ClpP and DRD2 are highly expressed in medulloblastoma patient samples

To assess whether imipridones are a potential therapeutic strategy for MB patients, we first analyzed the expression of the ONC206 molecular targets ClpP and DRD2 in MB patient samples at a transcriptional and protein level. In review of publicly available databases (R2: Medulloblastoma Genomics Analysis and Visualization Platform, https://hgserver1.amc.nl/cgi-bin/r2/main.cgi?dscope=MB&option=main and NCBI Gene Expression Omnibus (GEO) GEO2R; https://www.ncbi.nlm.nih.gov/geo/query/acc.cgi?acc=GSE85217), we found significantly higher mRNA levels of ClpP in MB tumors relative to normal brain and normal cerebellum (Figure 1A). Importantly, ClpP is highly overexpressed across all MB molecular subgroups (Figure 1B), suggesting ClpP may be a viable therapeutic target for MB treatment. In contrast, while the expression level of DRD2 in MB tumors is significantly higher relative to normal brain overall, it is more comparable to the level observed in normal cerebellum (Figure 1C), indicating that DRD2 may be more of a secondary rather than a primary therapeutic target for MB. We then analyzed a cohort of MB tumors (N=24) from the Pathology Department of Children’s Healthcare of Atlanta, and at the protein level, we found evidence for high levels of ClpP expressed in all MB tumors analyzed, including all major molecular subgroups (6 tumor samples/group), while ClpP protein was undetectable by IHC in normal brain and normal cerebellum (Figure 1D). To validate our ClpP protein expression findings in MB, we interrogated the proteomic data available within a public database of pediatric brain tumors (CPTAC Data Portal https://proteomics.cancer.gov/data-portal), and confirmed that among pediatric CNS tumors, embryonal tumors, including MB and atypical teratoid/rhabdoid tumors (ATRT), have the highest level of ClpP expression relative to normal brain (Figure 1E).

**Figure 1.**
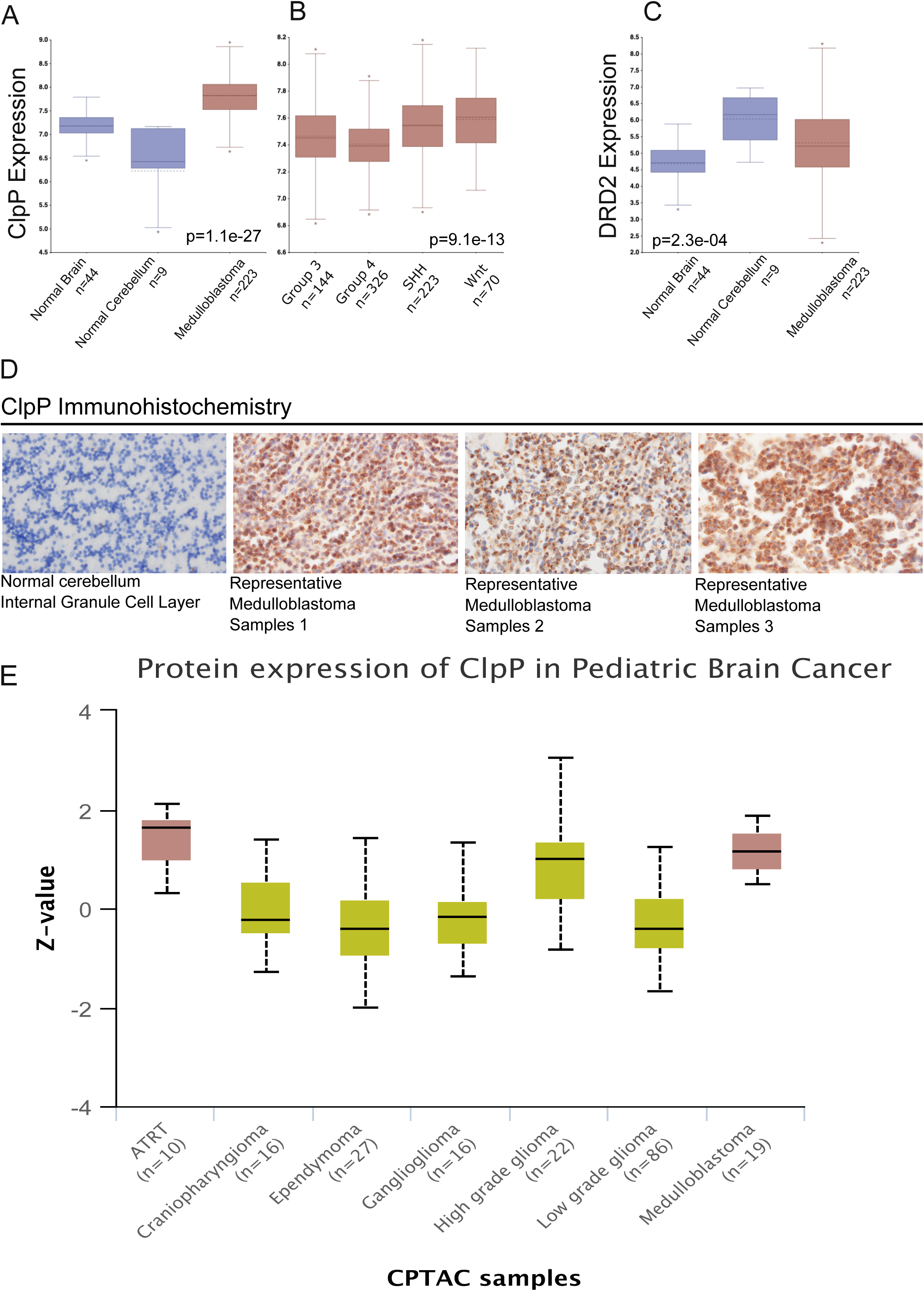
High expression of ClpP in primary medulloblastoma tumors A: RNA expression data from R2 public database showing high ClpP expression in 223 medulloblastoma tumors (MB) as compared to 44 normal brain (NB) and 9 normal cerebellum (CB) samples (p=1.1e-27). B: RNA expression data from R2 public database showing high ClpP expression in 763 medulloblastoma tumors across all molecular subgroups (p=9.14e-13). C: RNA expression data from R2 public database showing high DRD2 expression in 223 medulloblastoma tumors (MB) as compared to 44 normal brain (NB), yet at similar levels compared to 9 normal cerebellum (CB) samples (p=2.3e-4). D: Representative immunohistochemisty of tissues obtained from the Neuropathology Department of Children’s Healthcare of Atlanta showing low expression of ClpP in normal brain relative to medulloblastoma tumors with high protein expression. E: Plot of standard deviations from the median (z-value) across pediatric CNS tumor samples withing the CPTAC public database showing the highest protein expression levels for ClpP in embryonal tumors, ATRT and medulloblastoma.

### Imipridones impair cell viability and induce apoptosis in human and murine medulloblastoma

Next, we studied the effects of treatment of medulloblastoma cells with imipridones ONC201 and ONC206 *in vitro*. We saw a pronounced reduction in cell viability of human Group 3 and Group 4 and murine Group 3 medulloblastoma cells (Figure 2), with ONC206 exhibiting a much lower IC-50 as compared to ONC201 (Figure 2H, IC-50 range for ONC201: 170nM – 5μM; for ONC206: 14nM – 2μM). Notably, ONC206 effectively reduced viability of DRD2 negative MB cells UW228, Daoy and D556 (Figure 2G).

**Figure 2.**
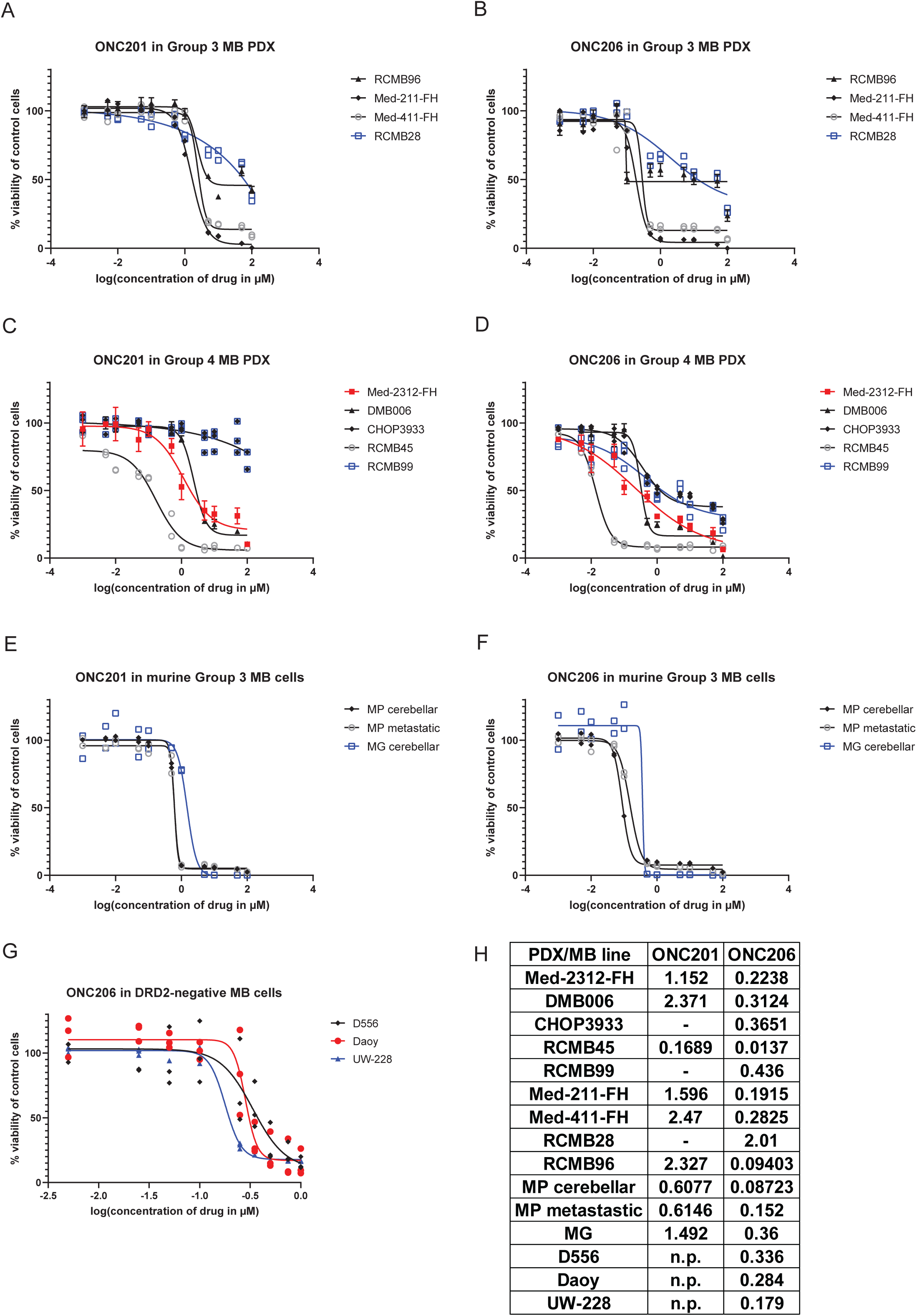
ONC201 and ONC206 reduce cell viability of human and murine medulloblastoma cells Cell viability assay of ONC201 (A, C, E) and ONC206 (B, D, F, G) treated medulloblastoma cells (human Group 3: A-B, human Group 4: C-D, murine Group 3: E-F, human DRD2-negative cells: G); H: IC-50s for each agent and medulloblastoma cell line.

### ONC201 and ONC206 induce mitochondrial damage and ATF4-dependent apoptosis in human MB cells

To further explore the molecular mode of action of imipridone treatment in medulloblastoma, we assessed protein changes after ONC201 and ONC206 treatment of human MB cells and noted that both agents induced mitochondrial damage. Following treatment of D556 cells with either drug, we observed an induction of mitochondrial hyperpolarization (Figure 3A and B) in concert with a marked downregulation of electron transport chain complexes and Survivin (Figure 3C and D). Notably, a relative increase of cytochrome c within the hyperpolarized mitochondria, indicating induction of early apoptosis and mitochondrial dysfunction, was seen upon treatment with ONC206 compared with ONC201 (Figure 3B), which may point to one mechanism resulting in greater anti-tumor potency of ONC206 compared to ONC201. We also observed a downregulation of ClpP in Daoy and to a lesser extent in D556 cells (Figure 3B). Moreover, we saw induction of the ATF4 pathway, thereby implying that both agents induce an integrated stress response (ISR) (Figure 4A).

**Figure 3.**
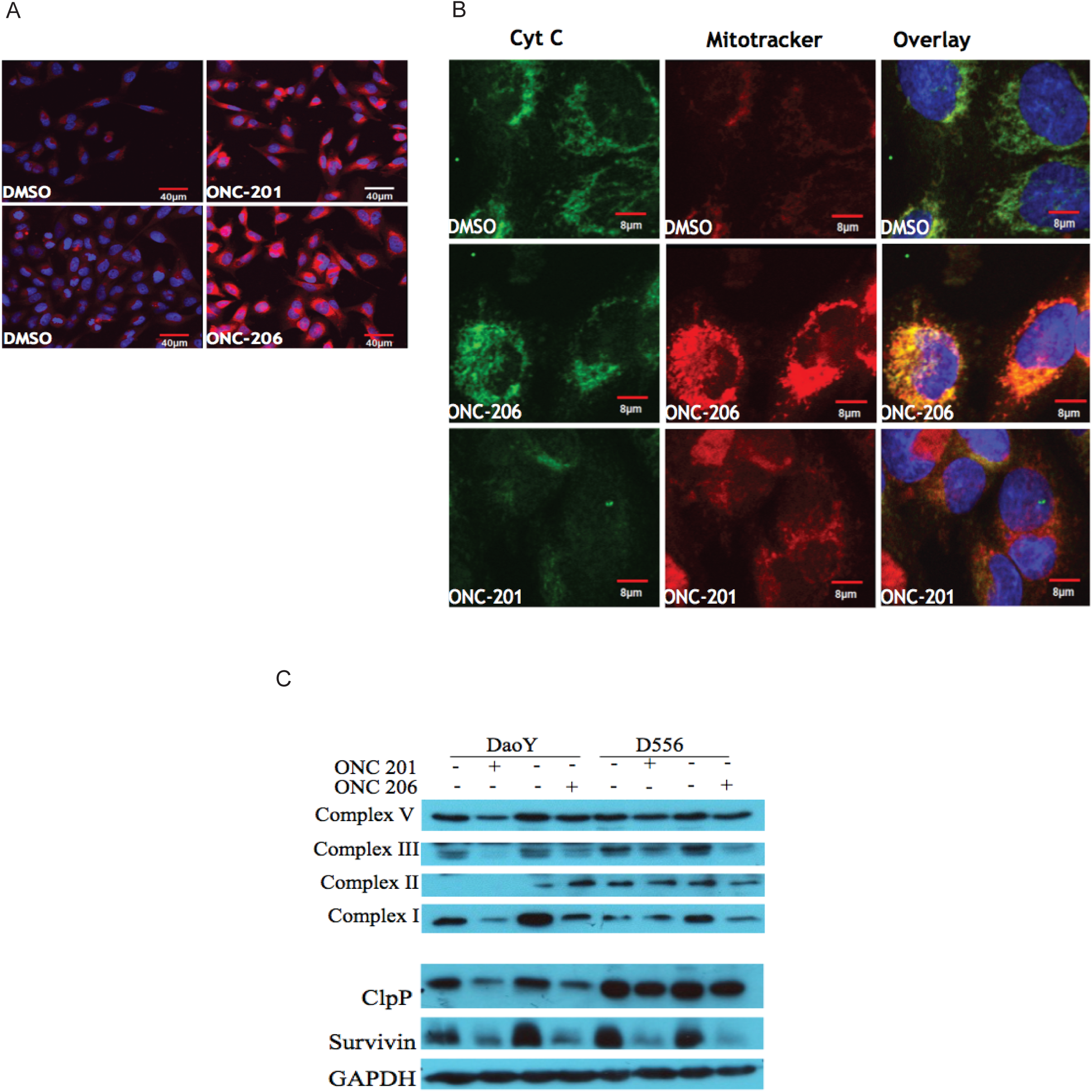
ONC201 and ONC206 induce mitochondrial damage in medulloblastoma cells A: Immunofluorescence staining with mitochondrial-specific cationic dye (MitoTracker Red) of D556 cells treated with ONC201, ONC206 or DMSO (overlay with DAPI) shows an accumulation of MitoTracker Red in ONC201/206-treated cells relative to the DMSO control-treated cells, indicating mitochondrial dysfunction as evidenced by a marked increase in mitochondrial hyperpolarization. B: Cytochrome C and MitoTracker staining of ONC201, ONC206 and DMSO treated D556 cells demonstrating higher cytochrome C within the hyperpolarized mitochondria upon treatment with ONC206 relative to treatment withONC201, indicating that the ONC206 treated cells are specifically undergoing early programmed cell death and exhibiting mitochondrial dysfunction. C: Western blot for mitochondrial complexes I/II/III/V after ONC201 and ONC206 treatment of DaoY and D556 cells demonstrating protein downregulation of all complexes after treatment with both drugs. D: Western blot for of ClpP and Survivin after ONC201 and ONC206 treatment of DaoY and D556 cells demonstrating protein downregulation of Survivin, and to a lesser extent ClpP, after treatment with both drugs.

**Figure 4.**
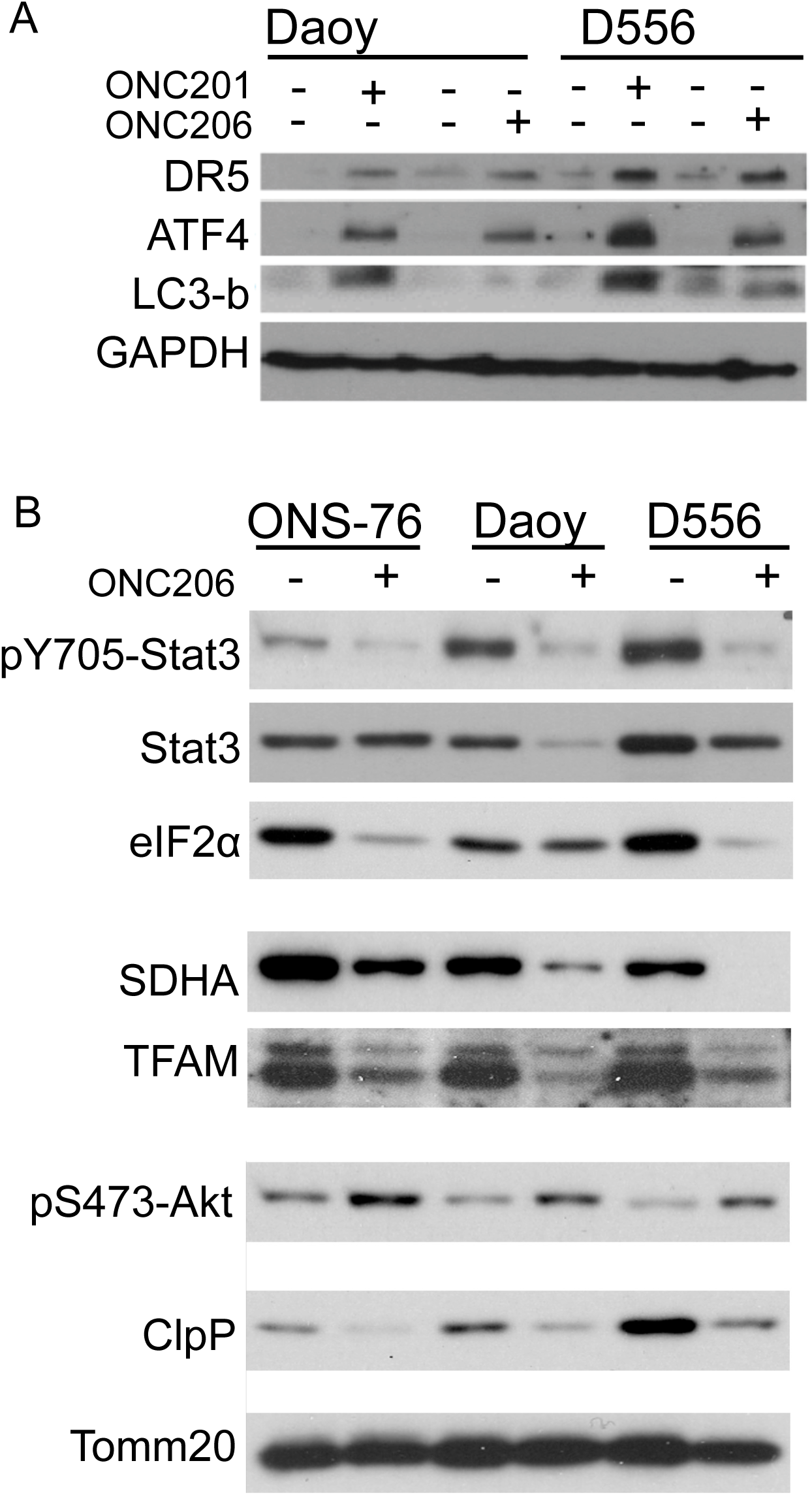
ONC201 and ONC206 induce ATF-dependent apoptosis, downregulation of distinct oncogenic pathways and upregulation of Akt A: Western blot for markers of ATF4 pathway demonstrating protein upregulation of DR5, ATF4, and LC3-b (ATF pathway activation) in DaoY and D556 medulloblastoma cells following treatment with ONC201 and ONC206. B: Western blot for oncogenic transcription factor STAT3 (total and activated pY705-STAT3), translation initiating factor eIF2a, and mitochondrial related proteins SDHA, TFAM, and ClpP demonstrating marked protein downregulation for each, with the exception of ClpP, following ONC201 and ONC206 treatment in ONS-76, DaoY and D556 medulloblastoma cells. phosphor-Akt (pS473) upregulation following treatment with ONC201 and ONC206.

To further elucidate the on-target effects of ONC206, we investigated the MB cells for changes in the protein levels of downstream mediators of mitochondrial function and cell survival following treatment. We saw a significant downregulation of mitochondrial related proteins (e.g. succinate dehydrogenase complex flavoprotein subunit A (SDHA) and transcription factor A, mitochondrial (TFAM)) and a reduction of key oncogenic factors such as Signal transducer and activator of transcription 3 (STAT3), activated STAT3 (phospho-STAT3) and eukaryotic translation initiation factor 2A (eIF2A, Figure 4B). Moreover, we observed a reduction of ClpP (Figure 4B) and an upregulation of activated Akt (phopho-Akt, Figure 4B), potentially indicating the induction of compensatory pro-survival mechanisms of the tumor cells in response to cell death and ClpP loss following treatment with ONC206.

### ONC206 prolongs survival in SHH-driven and Group 3 murine models, as well as Group 3 and Group 4 PDXs

Initially, we tested the potential of ONC206 to cross the blood-brain barrier and saw that ONC206 was not only able to reach the brain, but also enriched within the tumor (Supplementary Table1). Notably, the concentrations we measured were well above the IC-50 determined in our viability assays (highest IC-50 of 2μM corresponding to 0.8mg/L, thereby 0.8mg/KG -assuming a density of 1g/mL and therefore well below the lowest concentration we measured of 4.1mg/KG). In order to test the efficacy of ONC206 and potential biomarkers *in vivo*, we used two murine MB models and four human MB PDXs. Immunocompetent mice with genetically engineered SHH-driven MB (ND2:SmoA1 mice, in which an activated Smoothened allele is driven by the NeuroD2 promoter) were treated orally with 120mg/kg of ONC206 weekly for two weeks, and immunocompetent mice with orthotopically implanted syngeneic Group 3 MB cells harboring *Myc* overexpression and dominant negative *Trp53* (MP tumors) were treated weekly with 100mg/kg ONC206 for four weeks with doses selected based on safety tolerability for mice of each model. Two Group 3 (Med-211-FH and Med-411-FH) and two Group 4 (Med-2312-FH and RCMB99) human MB PDX models were also chosen to evaluate the efficacy of ONC206 treatment. The Group 3 MB and the Med-2312-FH Group 4-bearing mice were treated with 100mg/kg ONC206 twice weekly for 4 weeks. Based on updates from ongoing clinical trials on safety and efficacy evaluating ONC206 dosing regimens, the ONC206 dose/schedule was adjusted, and RCMB99-bearing mice were treated with 50mg/kg twice a day for three consecutive days per week until disease-related death.

To assess potential biomarkers of ONC206 response in MB, we studied changes in previously described ONC201 treatment-associated biomarkers by real-time qRT-PCR and immunohistochemistry. We found that ONC206 treatment resulted in downregulated mRNA levels of *MYC* and upregulated mRNA levels of *EZH2* (Enhancer of Zeste 2 Polycomb Repressive Complex 2 Subunit) relative to control tumors, despite the *EZH2* results not reaching statistical significance (Supplementary Figure 3A,B). We validated each of these biomarker changes at the protein level in addition to demonstrating increased protein expression of death receptor 5 (DR5), TGM2 (tissue transglutaminase 2) and the apoptosis marker Cleaved-caspase 3, and decreased protein expression of NDUFS7 (NADH Dehydrogenase [Ubiquinone] Iron-Sulfur Protein 7, borderline non-significant), and ClpP (Figure 5A) in ONC206-treated, as compared to control treated tumors.

**Figure 5.**
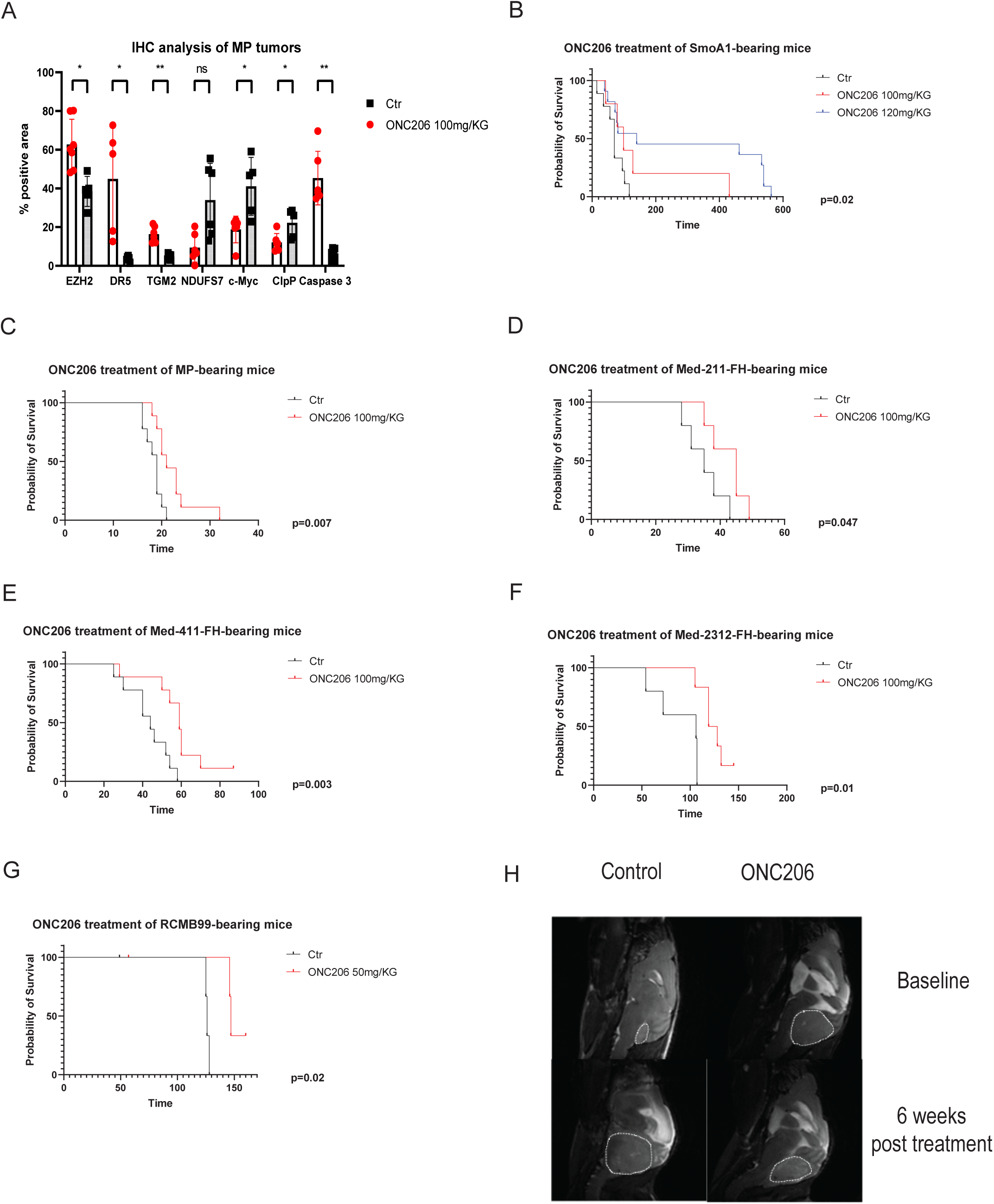
ONC206 prolongs survival in human and murine medulloblastoma A: IHC analysis of tumor markers for ONC206 on-target treatment effects in murine Group3 medulloblastoma bearing mice. The y axis represents the % of area positive for each individual marker showing an ONC206-induced upregulation of EZH2, DR5 and TGM2 and a downregulation of NDUFS7, c-Myc and ClpP relative to DMSO control-treated tumors. B-C: Survival of syngeneic mice with murine SHH-driven (B: SmoA1, prolongation of survival from 70 to either 99 days with 100mg/KG/week or 140 days with 120mg/KG/week of ONC206, p=0.02) or Group 3 (MP) medulloblastoma cells (C, prolongation of survival from 18 to 21 days with 100mg/KG twice weekly of ONC206, p=0.007). D-E: Survival of mice with Group 3 PDX medulloblastoma cells (D: Med-211-FH, prolongation of survival from 35 to 45 days with 100mg/KG twice weekly of ONC206, p=0.047; E: Med-411-FH: from 44 to 59 days with 100mg/KG ONC206 twice weekly, p=0.003 ) F-G: Survival of mice with Group 4 PDX medulloblastoma cells (F: Med-2312-FH, prolongation of survival from 106 to 124 days, p=0.01; G: RCMB99 from 126 to 147 days, with 50mg/KG twice daily three times a week of ONC206, p=0.02). H: Representative MRI images of ONC206 and DMSO control-treated transgenic mice with murine SHH-driven medulloblastoma (SmoA1) showing that tumor is significantly smaller after 6 weeks of treatment with ONC206 relative to baseline and compared to tumor at 6 weeks in the DMSO control-treated mouse.

In terms of efficacy *in vivo*, ONC206 led to a significant prolongation of survival in both murine MB models, with ND2:SmoA1 mice demonstrating a dramatic survival benefit when treated with this agent (prolongation of survival from 70 to 140 days, p=0.02; Figure 5B) and MP tumor-bearing mice showing a more modest effect (prolongation of survival from 18 to 21 days, p=0.007; Figure 5C). Notably, ONC206 significantly decreased tumor growth over time, as demonstrated by longitudinal magnetic resonance imaging (Figure 5H, Supplementary Figure 1A,B). While all MP mice (both ONC206 and vehicle treated) succumbed to their disease, ONC206 treated mice interestingly had smaller tumors at their endpoint as compared to control mice (Supplementary Figure 2A).

Mice from both Group 3 PDXs and the Group 4 PDX also responded to ONC206, with two of the eight Med-411-FH animals exhibiting long-term tumor-free survival (Med-211-FH: prolongation of survival from 35 to 45 days, p=0.047 (Figure 5D); Med-411-FH: from 44 to 59 days, p=0.003 (Figure 5E); Med-2312-FH: from 106 to 124 days, p=0.01 (Figure 5F); RCMB99 from 126 to 147 days, p=0.02 (Figure 5G)). Of note, ONC206 also led to significantly delayed tumor growth as compared to control tumors in RCMB99-bearing mice (Supplementary Figure 2B). We thus were able to show safety and efficacy of ONC206 treatment *in vivo* with long-term survivor mice in both the setting of murine, immunocompetent models and human PDX cells.

## Discussion

Our results demonstrate that ONC206 is an effective therapeutic that warrants further clinical investigation for the treatment of children with high-risk medulloblastoma. We have identified universal upregulation of ClpP, a key mediator of the effects of ONC206, in MB patient samples, and have shown that medulloblastoma cells respond to imipridones and particularly ONC206 *in vitro*. Moreover, we have identified mitochondrial disruption and ATF4-related apoptosis as the molecular mechanism of action for ONC206 in medulloblastoma cells. Finally, we have demonstrated CNS tumor drug penetrance, safety and efficacy of ONC206 treatment *in vivo* across diverse murine MB models, including immunocompetent models that recapitulate SHH and Group 3 MB and in human Group 3 and Group 4 MB PDXs.

Despite the initial assumption that the main mechanism of action of the imipridone class of drugs ONC201 and ONC206 is antagonism of DRD2, thereby mediating apoptosis of tumor cells via the integrated stress response, subsequent research has shown that these drugs act primarily via binding and activation of mitochondrial ClpP, thereby inducing mitochondrial damage,^12^ and that this anti-tumor mechanism is more potent than DRD2 targeting.^12,25^ We confirmed the ONC206-mediated mitochondrial damage and apoptotic-promoting on-target effects in DRD2 negative MB cells and showed that ClpP is ubiquitously expressed in murine and human MB tissues and cells, implying that the *in vitro* and *in vivo* effects of ONC206 in this study are the result of targeting ClpP. These results suggest that ClpP rather than DRD2 should be used as a primary biomarker in the clinical investigation of ONC206 for the treatment of patients with MB.

Przystal et al. have described the ability of ONC206 to induce mitochondrial alterations including an increase of reactive oxygen species (ROS) and a decrease in oxidative phosphorylation and ATP production thereby leading to apoptosis of DMG cells via interaction with ClpP ^25^. This metabolic reprogramming is of high importance, since cancer cells have been shown to energetically depend on mitochondrial function for their survival ^26,27^. Our results show a similar induction of mitochondrial damage, herein first described in MB, and are in line with current literature indicating that ONC206 is more potent in induction of these mitochondrial changes than ONC201 ^18,25^. Interestingly, we saw effects on mitochondrial disruption with both agents, yet observed cytochrome c release, which is one of the most significant markers of mitochondrial danger-associated molecular patterns ^28^ and mitochondrial cell death ^29^ only after treatment with ONC206.

While ONC206 has been reported to downregulate AKT in glioblastoma and other tumors ^30,31^, we have found that ONC206 induces AKT phosphorylation in MB similar to the effects of ONC201 in DMG tumors ^32^. Future studies will determine potential synergisms and combinatorial strategies for ONC206 in the setting of MB. Based on our results, AKT inhibitors (currently in clinical trials for various cancers, including brain tumors ^33,34^) could be interesting candidates for such an approach. Jackson et al. recently reported beneficial effects of combining ONC201 and the PI3K/AKT inhibitor Paxalisib, an agent that could also be considered as an add-on strategy for ONC206 ^32^. Alternatively, ONC206 could be combined with Poly ADP-ribose polymerase inhibitors, a strategy that has shown to downregulate AKT and produce overall promising results in endometrial cancer ^35^.

Our study and others have shown not only preclinical efficacy, but also safety for ONC206 in the treatment of brain tumors. Notably however, Przystal et al, who saw moderate responses in DMG to ONC206 *in vivo*, used much lower doses (50mg/KG, once weekly) than our study (100-120mg/kg twice weekly or 50mg/kg b.i.d. thrice weekly). This may be due to relative differences in blood-brain-barrier penetration between the models tested and/or target expression, including a higher prevalence of DRD2 on glioma cells. The pronounced survival benefit we observed (particularly in our SHH-driven model) might be attributed to the higher dose of ONC206, although differential responses of different tumor types are certainly conceivable. Other possibilities are that MB cells are more sensitive to the effects of mitochondrial damage or that in immune-competent murine models, such as our SHH MB model, drug-induced cell death may also induce a local immune reaction that augments the anti-tumor properties of the drug. Our study was limited in that we did not examine local changes in immune infiltration following treatment. Interestingly, a novel mass spectrometric assay developed to measure ONC206 in patient plasma showed that a dose of 50mg reached a plasma peak level (PPL) of about 250ng/mL, whereas a dose of 100mg led to one of more than 400ng/mL ^36^. Beyond the ONC206 dose, which is currently being tested in a dose-escalating trial for recurrent pediatric brain tumors (PNOC-023), our data are very encouraging, since even the lowest detected PPL of 250ng/mL corresponds to 612nM and is thus much higher than 11/12 IC-50s for ONC206 we detected in our metabolic viability assays with human and murine cells.

Biomarkers for treatment choice and response are of utmost importance in the era of targeted therapy, especially due to intertumoral heterogeneity. Previous studies have highlighted biomarkers of response to ONC206 ^37–39^. In our study, we were able to confirm changes in some of the known biomarkers of ONC206 in Group 3 medulloblastoma *in situ*, most notably the upregulation of DR5 and Caspase 3, indicating induction of apoptosis, and the downregulation of MYC, a key oncogene in Group 3 medulloblastoma. Interestingly, we saw an ONC206-induced increase of EZH2. Prior studies have shown ONC201 synergizes with EZH2 inhibitors ^40^. This could be considered as a compensatory mechanism of tumor cell survival to ONC206 treatment thereby potentially supporting a combinatorial treatment of ONC206 and EZH2 inhibitors. However, inactivation of EZH2 has been described to epigenetically induce expression of Gfi1 in MB, which then together with MYC supports tumor growth in Group 3 medulloblastoma ^41^. Therefore, our observed ONC206-associated upregulation of EZH2 could potentially be leading to histone repression of MYC and thereby downregulation of MYC levels, which we also observed after ONC206 treatment. Together, this points toward a potential alternative mechanism of action of ONC206 that may be unique to medulloblastoma. Future studies are warranted to unravel the complex interaction between EZH2 and MYC and potential effects of ONC206 in this context.

In summary, our results highlight ONC206 as a potentially attractive new therapeutic option for patients with treatment-resistant MB. Future studies are warranted to identify potential resistance mechanisms that may be overcome by combinatorial strategies for ONC206 in the treatment of MB. Importantly, our results have paved the way for a clinical trial (NCT04732065) testing the efficacy of ONC206 in recurrent malignant brain tumors, including medulloblastoma.

## Ethics Statement

All animal experiments were approved by the Institutional Animal Care and Use Committee of both research institutions (Emory University and Sanford Burnham Prebys Medical Discovery Institute).

## Funding

This work was supported by funding from The Cure Group 4 Consortium (TM and R.J.W-R), as well as funding from the German Research Foundation (DFG) to TT (Reference number: TZ 102/1-1; project number: 454298163), the ChadTough Defeat DIPG Foundation to TT, the National Institute for Neurological Disorders and Stroke (R35 NS122339 to RJW-R) and the National Institute on Aging (P01 AG073084 to PDA). SBP’s Shared Resources are supported by SBP’s NCI Cancer Center Support Grant P30 CA030199. TM’s and RJWR’s laboratories were also funded by Ian’s Friends Foundation and Alex’s Lemonade Stand Foundation. TM’s group was additionally funded by the Lassiter Family Foundation and the CURE Childhood Cancer, and RJWR’s group by the V Foundation, William’s Superhero Fund and the McDowell Charity Trust.

## Conflict of interest

VVP and JEA are employees of Chimerix and Jazz. VVP and JEA are shareholders of Chimerix, Oncoceutics and Jazz Pharmaceuticals. VVP and JEA are inventors on dordaviprone- and ONC206-related patents. All other authors have no interest to declare.

## Authorship

TT designed the study, conducted experiments and acquired and analyzed data. JL, FLC, AM, DZ, HZ, JEVV, MS, TS and PDA conducted experiments, provided reagents, analyzed data and helped with data interpretation. JEA and VVP provided reagents and along with RJWR analyzed and interpreted data and planned different steps of the study. TJM designed and supervised the study as well as experimental procedures and data analysis. TT and TJM wrote the manuscript with input from all co-authors.

## Data availability

All data that support the findings of our study are available from the corresponding author upon request.

## Acknowledgements

We gratefully acknowledge the Animal Facilities at Emory University, UCSD and SBP for help with animal care and husbandry. We gratefully thank Guillermina Garcia and Monica Sevilla from the histology core facility for their help with our experiments.

The authors also acknowledge the funding agencies listed above.

---

Supplementary Table 1

ONC206 concentration in murine tumor and brain tissues measured by HPLC; SHH-driven SmoA1-bearing mice were given 120mg/KG of ONC206 (n=3) or vehicle control (n=3), mice were euthanized one hour later and tumor and non-tumor surrounding normal cerebellar tissue were collected, followed by HPLC analysis. ONC206 was detected in the surrounding cerebellum and at higher concentrations within the tumor and not detected in the vehicle treated murine tumor and cerebellar tissues (bql = below quantification limit). Depicted are the values from individual mice, as well as the mean and standard deviations.

Supplementary Figure 1

MRI-measured tumor volumes (mm3) of SHH-driven SmoA1 transgenic mice treated with either 120mg/KG or 100mg/KG ONC206 or vehicle control. Depicted are volumes at baseline, demonstrating no significant difference in tumor volumes at the start of treatment between groups, and 12 weeks after treatment initiation, demonstrating significant tumor reduction with ONC206 treatment in a dose-dependent manner and compared to control-treated tumors.

Supplementary Figure 2

A: Tumor cell yield from Group 3 (MP)-bearing syngeneic mice at survival end-point; Mice were sacrificed and tumors were dissociated according to our described protocols. At the end of the dissociation, the number of viable cells was measured and the total number of cells per tumor (n=4 from ONC206-treated and n=4 from vehicle-treated mice) is depicted in the y axis. Data demonstrate significantly lower tumor cell yield following ONC206 treatment compared to control treated tumor.

B: Longitudinal IVIS data for RC99-bearing mice; At d40 after transplant, mice were randomized into two groups (n=4 per group) and were imaged at different time-points. Depicted is the IVIS signal in photons/second.

Supplementary Figure 3

qRT-PCR results of tumors from Group 3 (MP)-bearing syngeneic mice at survival end-point (n=4 from ONC206-treated and n=4 from vehicle-treated mice); depicted is the normalized expression ratio (A: MYC/β-Actin; B: EZH2/β-Actin) in control versus ONC206-treated tumors, demonstrating a relative decrease in MYC and increase in EZH2 following ONC206 treatment compared to control treatment (*p=0.02).

