## Supplementary Material for "ONC206 demonstrates potent anti-tumorigenic activity and is a potential novel therapeutic strategy for high-risk medulloblastoma"

### Slide 1
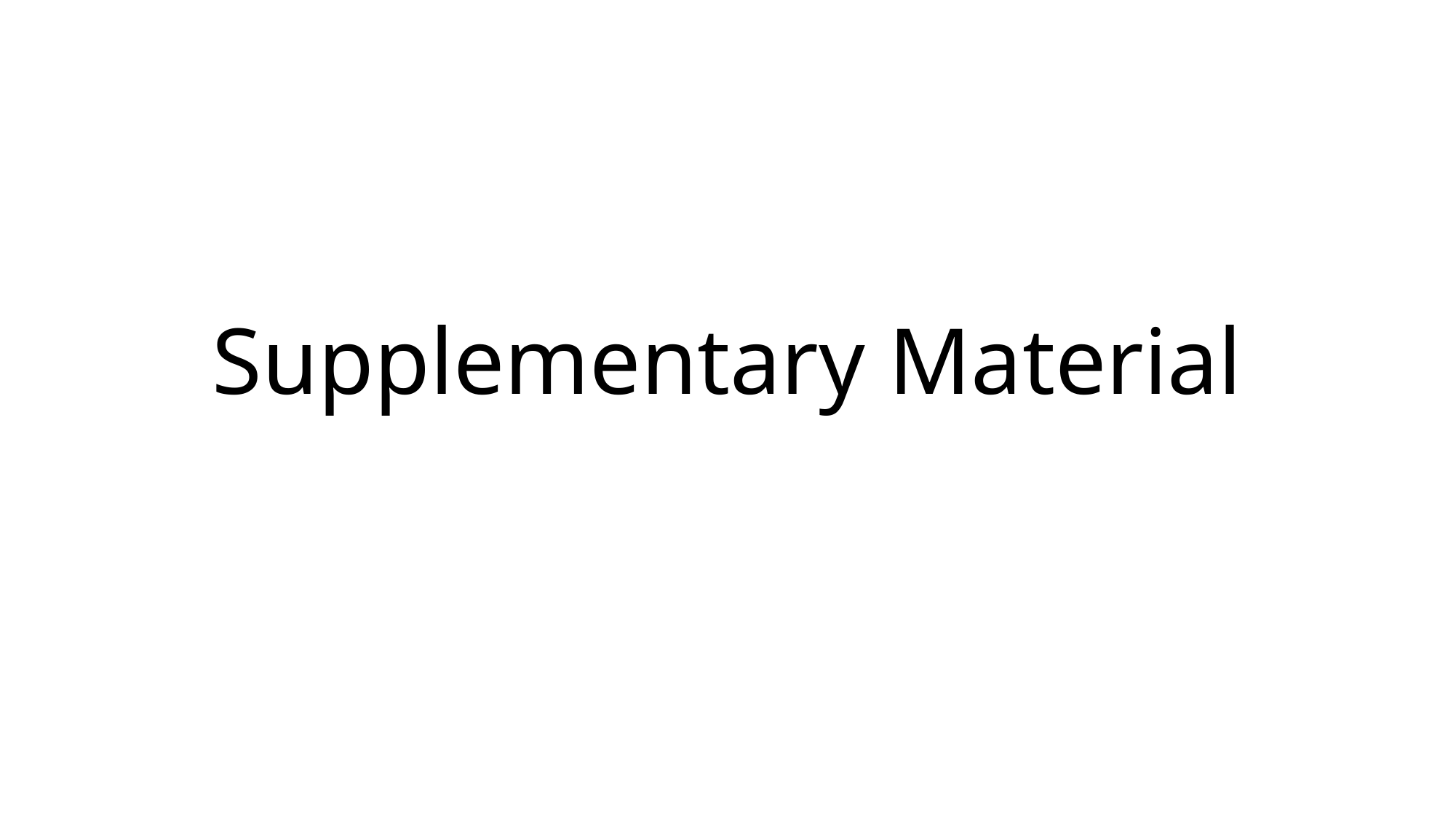

#### Supplementary Material

### Slide 2
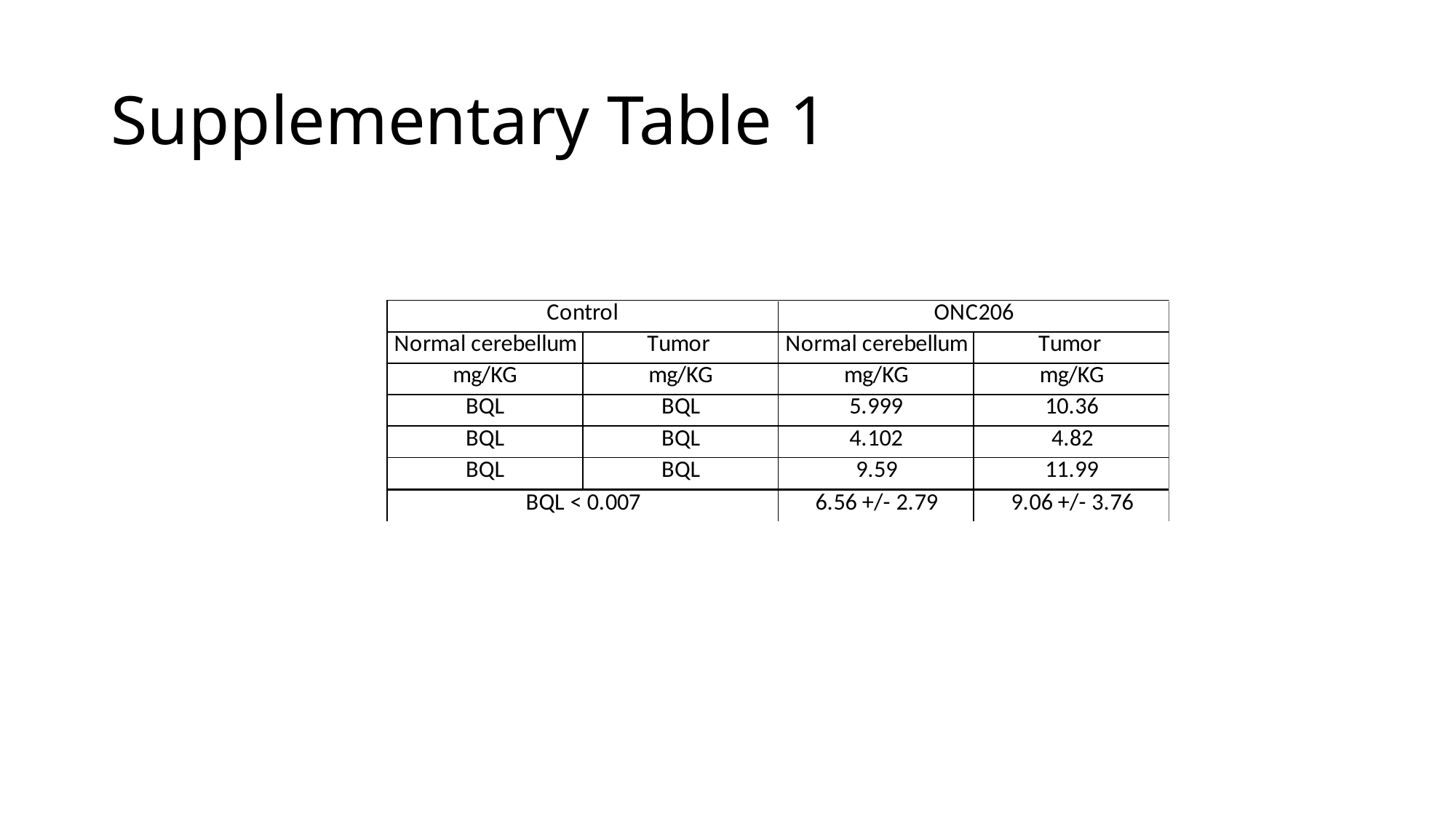

#### Supplementary Table 1

### Slide 3
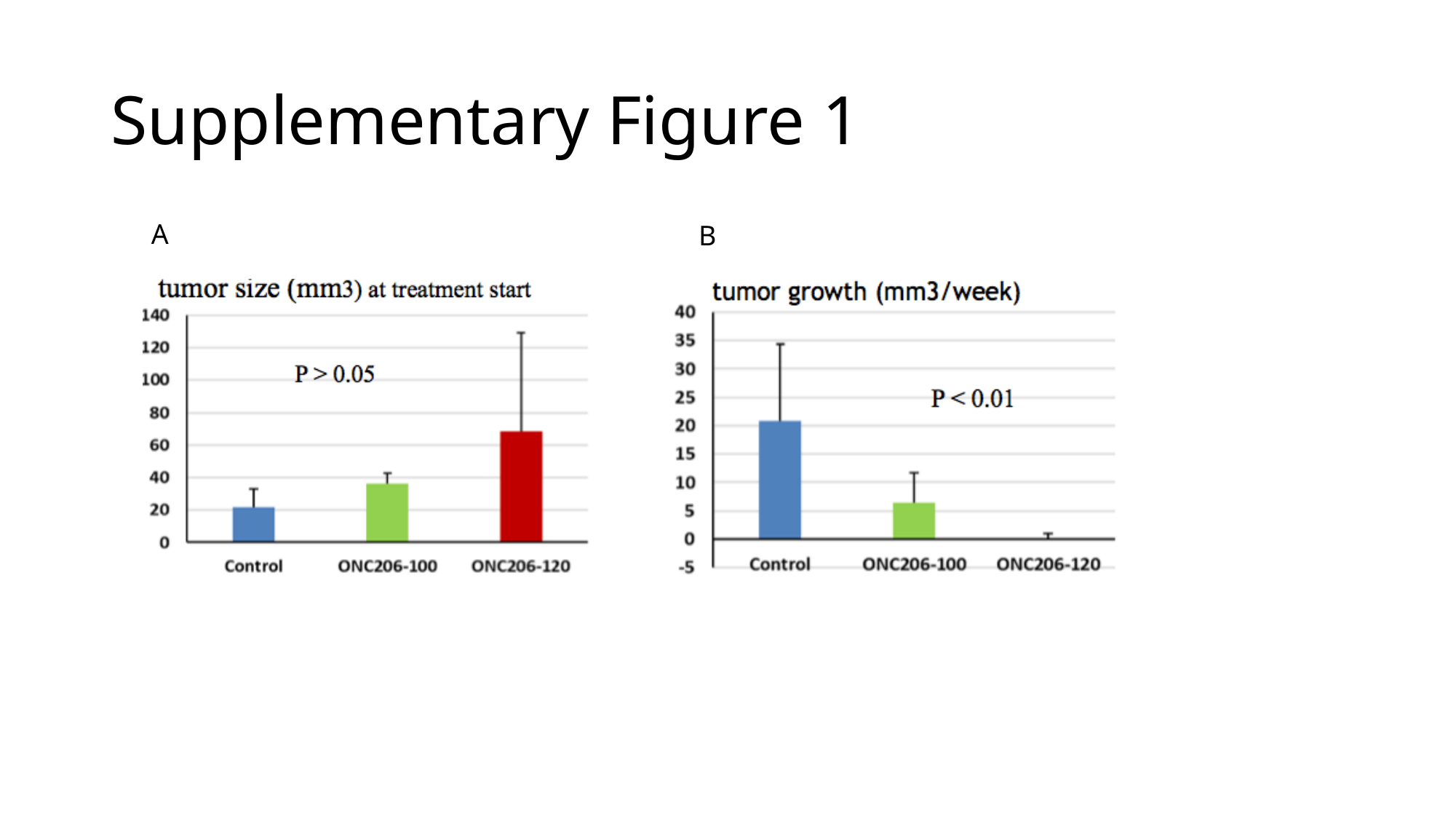

#### Supplementary Figure 1
A
B

### Slide 4
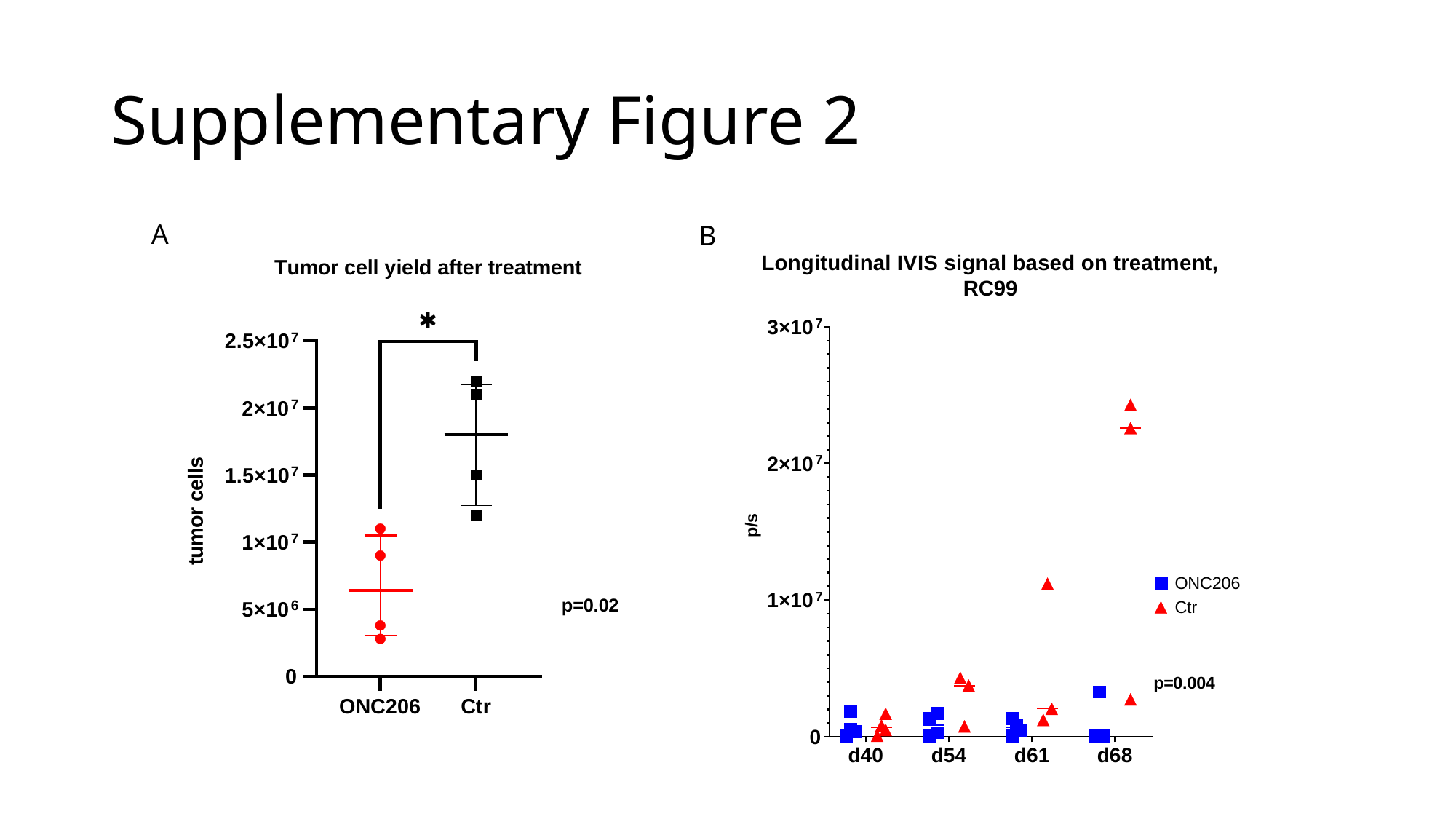

#### Supplementary Figure 2
A
B

### Slide 5
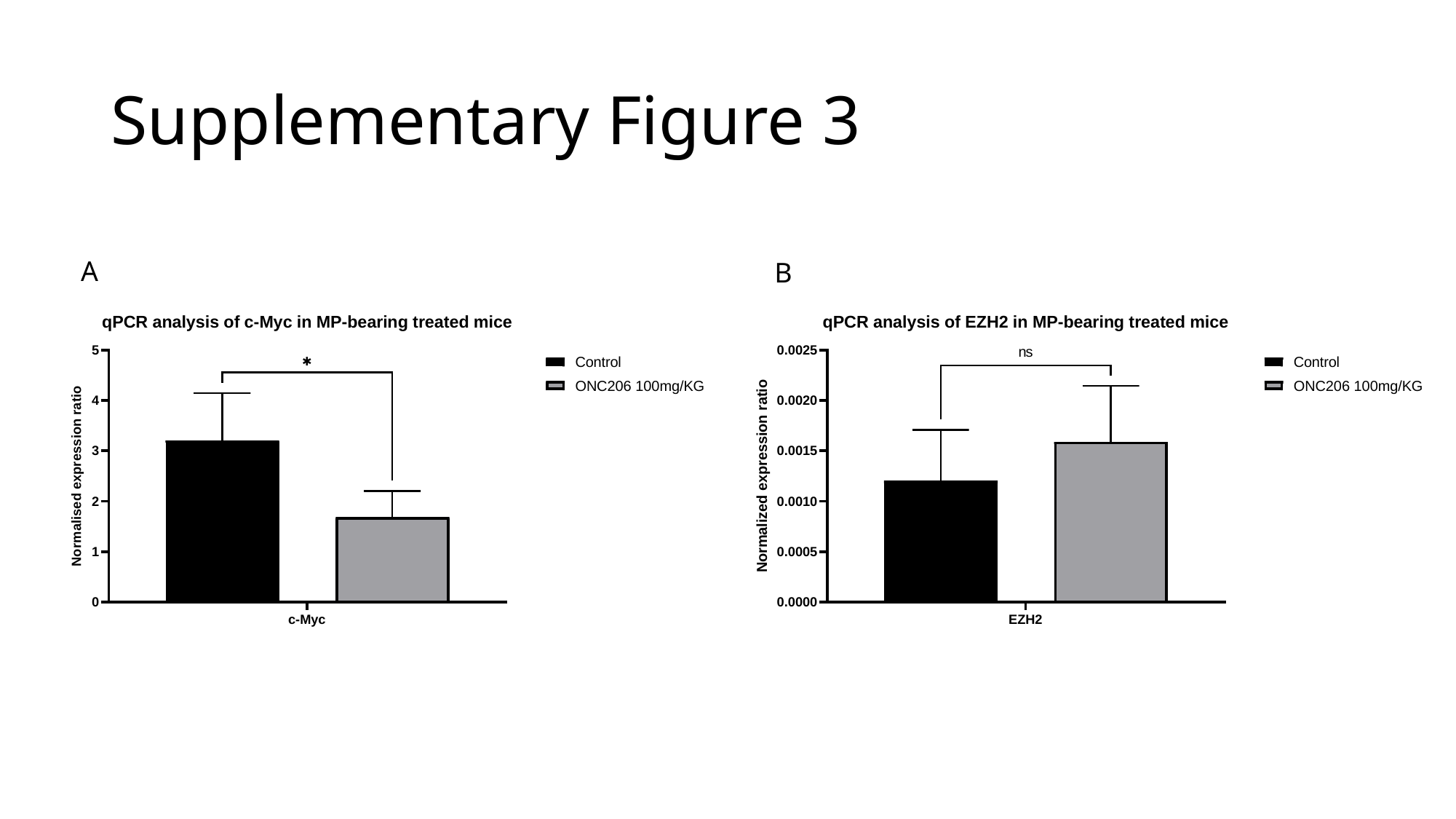

#### Supplementary Figure 3
A
B
